# Common Neural Code for Reward and Information Value

**DOI:** 10.1101/324665

**Authors:** Kenji Kobayashi, Ming Hsu

## Abstract

Adaptive acquisition of information is critical for goal-directed behavior. Popular theories posit that information acquisition is driven by intrinsic motives (curiosity or exploration bonus) and mediated by valuation system. However, they are insufficient when agents need to evaluate instrumental benefit of new information in a forward-looking manner. We tested whether human brain computes value of information (VOI) on a scale common with more basic rewards to acquire information. In an fMRI task, subjects purchased information for choices on monetary lotteries. Behaviorally, subjective VOI was largely driven by instrumental benefit, as normatively predicted, but additionally affected by non-instrumental motive, particularly the utility of anticipation. Neurally, VOI was represented in striatum, ventromedial prefrontal cortex, and dorsolateral prefrontal cortex. Cross-categorical decoding revealed that these regions use a common scale for VOI and another type of value, expected utility of the lotteries. These provide new insight on neurocognitive mechanism of forward-looking, value-based information acquisition.

## Introduction

Adaptive acquisition of information is critical in goal-directed behavior in humans. Collecting too little information, paying too much for information, not discriminating relevant information from irrelevant one, or acting on unreliable or false information, can all result in failure to achieve desired goals. Understanding neurocognitive mechanisms of adaptive information acquisition is not only important in neuroscience, psychology, and economics, but also has wide real-world applications, such as policymaking, public health, and artificial intelligence.

Information seeking behavior is frequently viewed as reflecting agents’ curiosity, i.e., motive to collect information for its own sake^1–4^. Explaining curiosity is a challenge to decision-making models, such as reinforcement learning (RL), because it is not directly reinforced by explicit, tangible rewards. To incorporate curiosity-driven information acquisition, decision-making models often postulate that acquisition of information is intrinsically rewarding, and more specifically, exploratory actions (or, in general, actions that would not be extrinsically rewarded) are encouraged by some forms of bonus utility^5–7^. Various forms of utility bonus have been proposed, such as surprise^8^, novelty^9–11^, perceived information gap^2^, and anticipatory utility (savoring and dread)^12–14^. At the neural level, dopaminergic reward system may multiplex intrinsic bonus utility with signals on extrinsic reward^15–17^. Multiplexing extrinsic and intrinsic rewards would make otherwise myopic agents to explore the environment and gather information, which may lead to improvement of decision making in the long term.

The idea that information acquisition is solely driven by curiosity or utility bonus, however, has been challenged on conceptual and empirical grounds. Most importantly, agents should be more motivated to acquire information if it has larger instrumental benefits under the current goal. For instance, we are interested in weather forecast if we are trying to decide whether to go hiking or reading indoors, but not so much if we have already decided to stay indoors. Such goal-driven information acquisition is particularly challenging when agents need to acquire information that they have never acquired before (e.g., a morning TV show in a foreign country we have never seen), where the bonus utility may not be adaptively formed based on the past history.

To evaluate information’s instrumental benefits in these cases, agents normatively need to be forward-looking and simulate their own actions and possible outcomes under different contents of information (“I’ll go hiking if it will be sunny, but reading indoors if rainy”). If agents are driven solely by curiosity but do not explicitly evaluate instrumental benefits, they may fail to discern relevant and useful information from irrelevant and useless one, which is problematic especially when information is costly. At the neural level, aforementioned curiosity-related dopaminergic activity is not sufficient for evaluation of instrumental benefits. It thus remains an open question to what extent goal-directed, forward-looking information acquisition is mediated by neural systems related to other types of reward and valuation.

The importance of instrumental benefit evaluation has been long recognized in economic and ethological studies of decision-making, owing to abundance of information acquisition in problems ranging from comparison shopping to job/mate search^18–20^. Standard economic accounts presume that agents are forward-looking and acquire information as a consequence of reward maximization. Information is acquired only if its cost is outweighed by value of information (VOI), i.e., how much the information would improve their choices and increases overall expected utility (EU). Although VOI calculation may be computationally more complex than more basic rewards (e.g., food or money), subsequent processes of cost-benefit analysis and action selection can be similar to other types of value-driven choices.

That the motivation to acquire information may be index by a single value measure opens up a number interesting possible hypotheses. First, dopamine reward system may drive information acquisition not only by encoding simplistic utility bonus but also by explicitly representing instrumental benefit. More precisely, while information acquisition may be driven by the combination of instrumental benefit and some non-instrumental motives (curiosity and/or utility bonus), valuation signals may multiplex these factors with each other and with other types of rewards. If this is the case, we would expect that VOI is represented on a neural common scale with other, more basic rewards^21^. Neural common currency is particularly advantageous when brains need to compare information acquisition actions against alternatives on the basis of their action values (i.e., exploration-exploitation dilemma)^5,22^. Although neural common currency has been examined in humans and monkeys^21,23–26^, it has never been tested with VOI, particularly when information is evaluated in the forward-looking manner.

To examine whether VOI is represented on neural common currency, we conducted an fMRI study where subjects made choices on costly, but directly actionable, information. Subjects were presented with a lottery with two monetary outcomes (a gain and a loss) and asked to choose whether to accept or reject it. The outcome probability was initially hidden and described as fair, but subjects could purchase the information to reveal the true probability. This information has direct instrumental benefit because subjects could change their choice flexibly based on the revealed probability. For instance, a subject may play a fair lottery with a large gain and a small loss, but reject it if the loss turns out to be more likely. Although there is a chance that the loss probability turns out to be smaller and she retains her original choice, she may purchase the information if the benefit of avoiding the loss is large enough to justify the cost.

We observed that subjects’ information acquisition behavior was indeed largely driven by instrumental benefit. Subjects’ information purchase choice was systematically sensitive to lotteries’ outcomes and possible probabilities, consistently with the standard VOI prediction. We further examined the contribution of additional non-instrumental motives. While we found no evidence for simplistic constant utility bonus, the utility of anticipation improved behavioral modeling. Next, using support vector regression (SVR) on voxel-wise BOLD signals, we found that the composite VOI was represented in traditional valuation regions, striatum and ventromedial prefrontal cortex (VMPFC). Importantly, cross-categorical decoding revealed that these representations were on a common scale with more basic values.

## Results

### Information acquisition is sensitive to instrumental benefits

To characterize the extent to which human information acquisition is sensitive to instrumental benefits, we used a two-stage task (Fig. 1a). Subjects were first asked whether to accept or reject a lottery with two outcomes (*x*_1_ and *x*_2_), assuming they would not receive further information (under the initial belief *s*_0_: *P*(*x*_1_) = *P*(*x*_2_) = 0.5). Next, two possible probability distributions were presented, *s*_1_ (*P*(*x*_1_) = π) or *s*_2_ (*P*(*x*_1_) = 1 − π), one of which would be true but revealed only if subjects purchased the information (Fig. 1b). π was manipulated on a trial-by-trial basis, either 2/3, 5/6, or 1. It determined diagnosticity of the information; it would perfectly predict the outcome if π = 1, but not much if π = 2/3. Subjects were then presented with the monetary cost of the information and indicated whether they would purchase it. Even though the true probability (*s*_1_ or *s*_2_) was not revealed during the task to prevent learning over trials, subjects were instructed beforehand that they would receive the information and could change their original choices after the scanning.

**Fig. 1.**
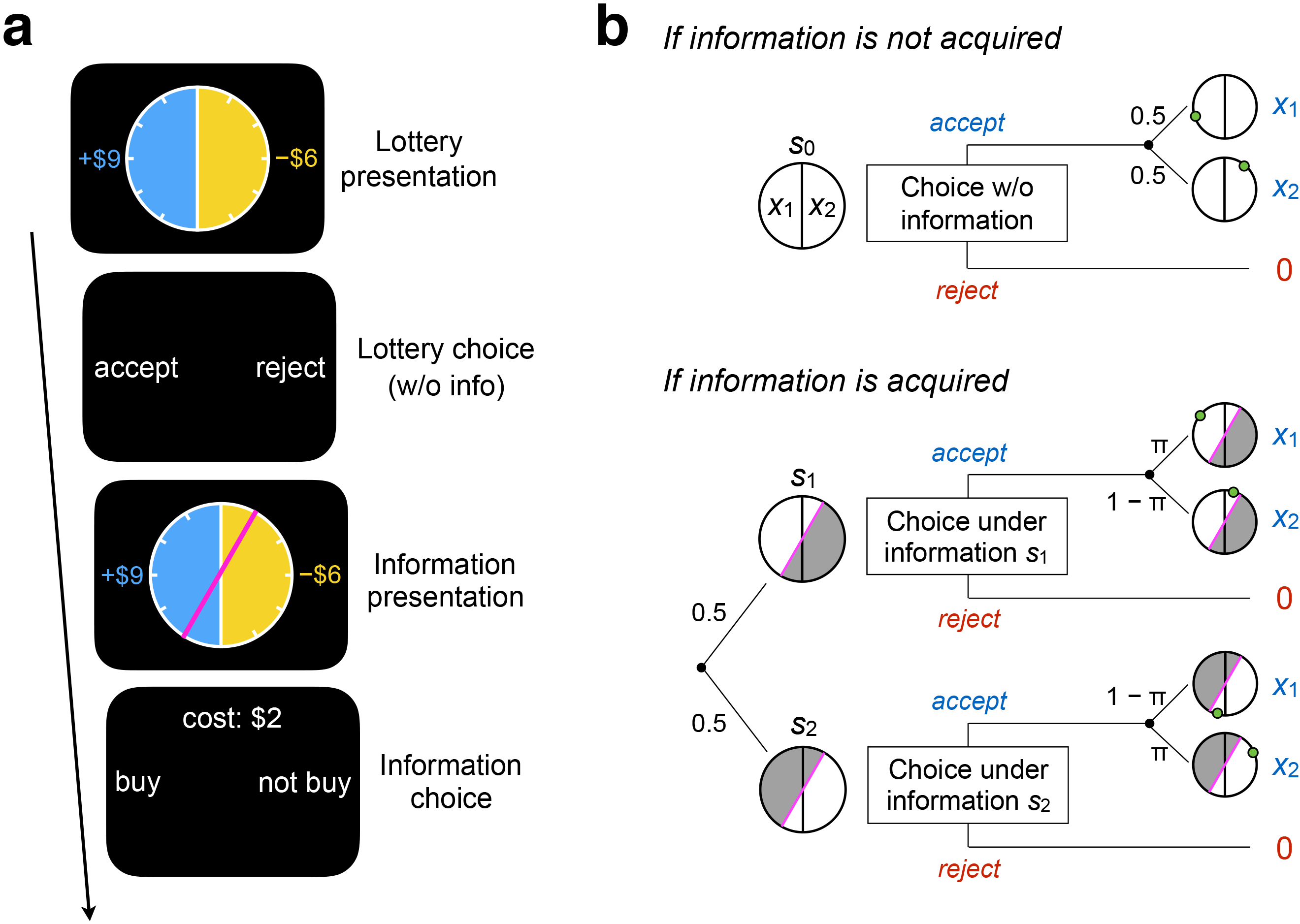
Experimental task design. (**a**) Subjects were presented with lotteries with two monetary outcomes, gain and loss, shown as a roulette wheel. When played, a green dot appeared at a random location on the perimeter, and its side determined the outcome (left or right). Outcomes were initially described as equally likely. Subjects indicated whether to accept or reject it, assuming they would not receive any further information. Potential information was then presented as a magenta partition. When purchased, it would reveal which side of the partition the green dot would appear. Subjects indicated whether to purchase it or forgo it given the information cost. Prior to scanning, subjects were informed that they could use purchased information to change their choices on lottery later. (**b**) *Top*: if subjects did not purchase information, they chose whether to accept or reject under the initial assumption (*s*_0_). *Bottom*: if subjects purchased information, it revealed that one of the two possible probability distributions, *s*_1_ or *s*_2_, was true. They were characterized by information diagnosticity, π, corresponding to the angle of the partition. Because subjects could not predict the true probability in advance, they need to stimulate *s*_1_ and *s*_2_ and average their EUs in order to compute instrumental benefit.

Under standard economic accounts, agents accept the lottery if its EU is higher than the utility of status quo *u*(0), and reject otherwise (Fig. 2a). Furthermore, they purchase the information if its cost is lower than VOI and forgo if higher. The information improves the overall expected utility only if their choices under *s*_1_ or *s*_2_ would differ from their choice under *s*_0_ (see Methods). VOI captures this marginal improvement of expected utility, i.e., the difference in the expected utilities between the decision with the information and the decision without. Note that, because agents cannot predict the true probability *a priori*, they need to simulate her own choices under *s*_1_ and *s*_2_, average their EUs, and compare it against EU under *s*_0_.

**Fig. 2.**
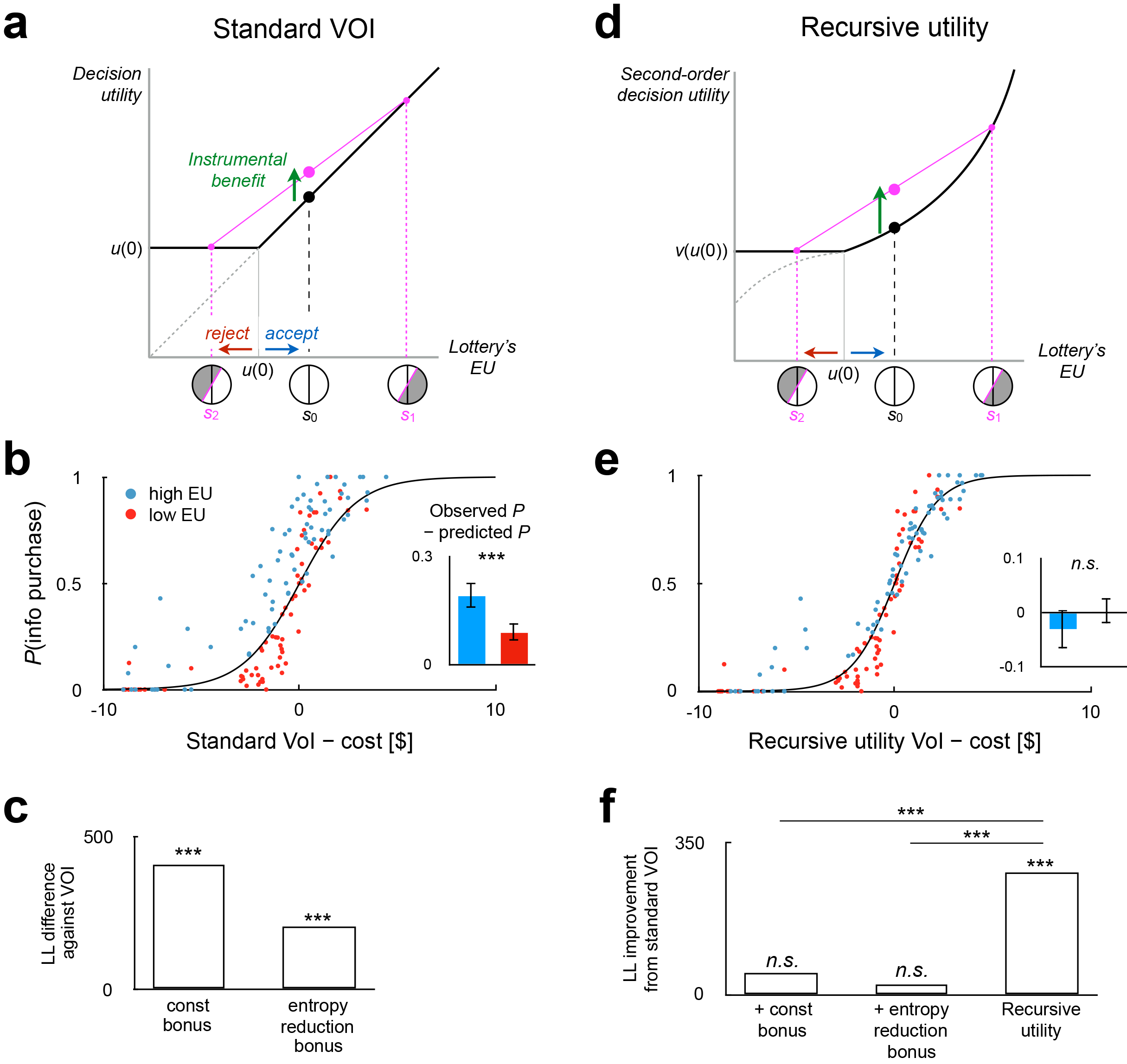
Behavioral results. (**a**) *Left*: Under standard VOI prediction, instrumental benefit is captured by the difference between the average EU of informed choices (*s*_1_, *s*_2_) and EU of uninformed choice (*s*_0_). Deterministic EU-maximizer is assumed, i.e., the lottery is accepted if its EU exceeded *u*(0). Instrumental benefit is positive as far as EU-maximizing choices differ under *s*_1_ and *s*_2_. *Right*: In the generalized VOI, instrumental benefit is modulated by anticipatory utility. This modulation is explained by non-linear second-order utility and may depend on stakes. (**b**) *Left*: subjects’ information purchase was sensitive to the difference between the instrumental-only VOI and cost. Inset: they exhibited over-purchase, particularly in high-EU lotteries (*blue*) more than low-EU lotteries (*red*). *Right*: the generalized VOI predicted information purchase better. Inset: stake-dependence in over-purchase is no longer apparent. Each dot corresponds to a unique combination of lottery, diagnosticity level, and cost, averaged over 37 subjects. Solid line curve: soft-max fit. (**c**) The instrumental-only VOI model achieved better goodness-of-fit (log likelihood, summed over 37 subjects) than utility bonus accounts. (**d**) The generalized VOI achieved better model fit than the instrumental-only VOI, while the composite model of the instrumental VOI and utility bonus did not. ***: *p* < .001, n.s.: *p* > 05.

Unlike many other hypothesized motivations of information acquisition, VOI computed as such is strongly sensitive to outcomes; VOI is large if both the potential gain and loss are large, and it is small if the potential gain is very large and the loss is trivial, or vice versa, because the agent would not change its choice irrespective of the true probability in the latter cases. We numerically derived the normative, instrumental VOI predictions based on outcomes (*x*_1_, *x*_2_) and diagnosticity (π), which predict subjects’ information purchase choices on a trial-by-trial basis. We found that the instrumental VOI was able to explain a substantial portion of the variation in information acquisition choices (evaluated based on binary choice likelihood, *p* < .0005 based on permutation of lotteries and diagnosticity).

We compared the normative instrumental VOI’s predictive power against two popular non-forward looking, non-instrumental motives: (i) a constant utility bonus, i.e., some fixed utility for novel information^5^, and (ii) a utility bonus scaled by entropy reduction, which is sensitive to π but still not to outcomes^2,6,7^. We found that VOI provided drastically better model fit (10-times 10-fold cross validation across participants, *p* < 10^−3^; Fig. 2c). This shows that subjects purchased information based on its instrumental benefit, and in particular the magnitudes of future possible outcomes, as normatively predicted.

### Coexistence of instrumental and non-instrumental motives

Although we found that the instrumental benefit is the main driver of information purchase in our task, some non-instrumental motive might additionally contribute. We next tested whether adding non-instrumental motives would improve the behavioral model fit. We specifically tested three models of non-instrumental motives from the literature: the two aforementioned utility bonus accounts (constant and entropy reduction) and anticipatory utility. Anticipatory utility, also often called “savoring” and “dread,” has been used in economics to justify people’s non-normative preference for information, and in particular timing of information delivery (e.g., many prefer to know if they win a raffle prize earlier because of savoring, while they prefer not to know the results of their cancer diagnosis because of dread)^12,14,27–30^.

We incorporated anticipatory utility in VOI calculations using recursive utility theory^12,31^. Recursive utility is similar to the standard VOI theory in that it assumes forward-looking and utility-maximizing agents. However, it allows the mere presence of information to increase or decrease the overall utility.

Under this assumption, therefore, agents may seek for information, not only because it has some instrumental benefit, but also because it improves the overall utility merely due to the difference in informational state. Specifically, the theory evaluates the lotteries in our task based on the expectation of second-order utility, which aggregates first-order EUs under the possible informational states in a nonlinear manner (Fig. 2a). If the aggregator function is convex, the difference in the overall expected second-order utility between choices with and without the information is amplified compared to the standard prediction, i.e., higher VOI. Conversely, if the aggregator function is concave, the marginal benefit in the expected second-order utility due to information is reduced (or sometimes reversed), leading to smaller VOI than the standard prediction. Therefore, recursive utility yields VOI that is the composite of instrumental benefit and anticipatory utility.

Importantly, since the non-instrumental component in recursive utility theory depends on the convexity of the aggregator function, it is naturally allowed to be dependent on the possible outcomes. This nicely echoes the intuitive general notion that savoring and dread tend to grow with the magnitude of the anticipated reward and punishment. For our purpose, the outcome-dependence of non-instrumental component is an important difference between recursive utility prediction and the other non-instrumental utility bonus accounts we deployed.

We found that our subjects’ behavior was consistent with this generalized VOI composed of instrumental benefit and anticipatory utility. It explained the observed information purchase better than the instrumental-only VOI (*p* < .001; Fig. 2b, d). On the other hand, neither of the utility bonus accounts improved the instrumental-only VOI model (*p* > .30). The difference in goodness-of-fit between the generalized VOI model and the utility bonus models was also significant (*p* < .001, respectively). These support the notion that non-instrumental motive is also sensitive to outcomes. Indeed, we noticed that subjects exhibited over-purchase of information, particularly among the lotteries with higher EUs (Fig. 2b left), which disappears when the behavior was compared against the anticipatory utility predictions (Fig. 2b right).

### Neural representation of VOI

The above results suggest that participants acquire information based on VOI, which is calculated in the forward-looking manner and combines instrumental benefit and anticipatory utility. We next sought to investigate the neural representation of VOI and its relationship with more basic reward values. In particular, we asked whether the VOI was represented in traditional value regions, and if so, whether that representation employs neural common currency. To this and, we asked subjects to make two types of value-based choices: whether to gamble on a lottery, and whether to acquire information regarding the said lottery. This allowed us to compare two types of value representations; the lottery’s EU, which is already widely studied, and VOI, which is the novel contribution of the current study.

We first looked for VOI representation during the presentation of the potential information’s diagnosticity. Combined with potential outcomes, which were already presented when gamble was first presented, the diagnosticity is sufficient for participants to compute subjective benefit of the information, both instrumental and non-instrumental components (Fig. 1a). Trial-by-trial numerical predictions of the generalized VOI was obtained from behavioral fitting of the recursive utility model described above. For voxel-wise BOLD signals in each searchlight (8mm radius), we conducted one-run-leave-out five-hold cross-validation using support vector regression (SVR) (Fig. 3; see Methods for details). Prediction accuracy was measured as correlation between the predicted and actual VOIs. In evaluation of correlation, we controlled for information diagnosticity π (i.e., partial correlation). This is to ensure that we detect regions engaged in valuation, rather than information-theoretic processing (e.g., responses to entropy reduction) or visual processing (π was represented saliently as the angle of the magenta partition; see Fig. 1a).

**Fig. 3.**
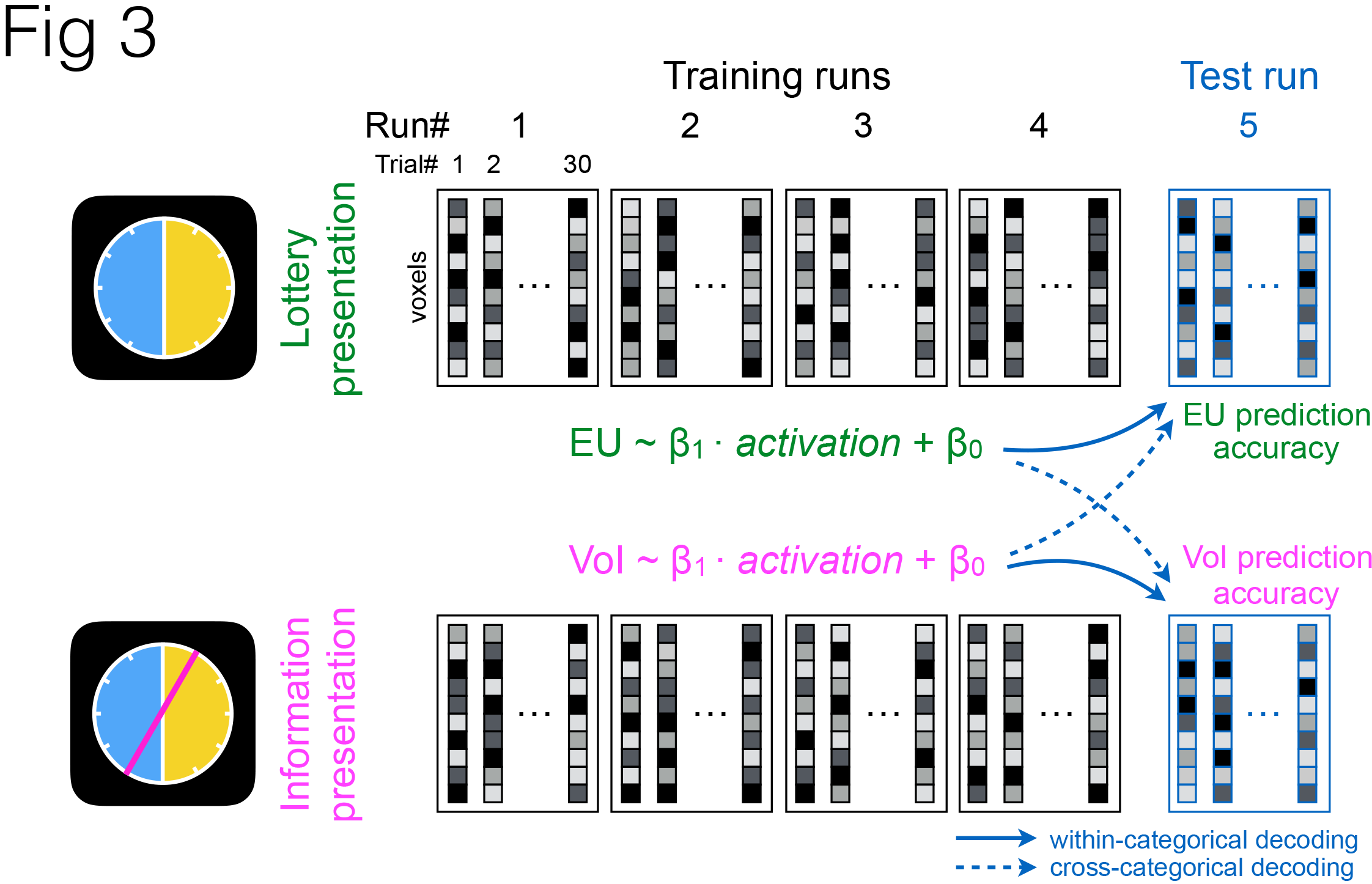
Schematic illustration of decoding analysis. Values were decoded from voxel-wise, trial-wise activation in two epochs, lottery presentation (top) and information presentation (bottom), using support vector regression (SVR) in searchlight. Decoder was trained using four runs and tested in the hold-out run (five-fold leave-one-run-out cross validation). In addition to the traditional (“within-categorical”) decoding approach (solid arrows), cross-categorical decoding (dotted arrows) was conducted to test the common currency hypothesis; decoder was trained on EU and tested on its predictability of VOI, and vice versa.

Consistent with the idea that dopaminergic reward systems are involved in value-driven information acquisition, we found that VOI was decodable from striatum and ventromedial prefrontal cortex (VMPFC) (*p* < .05, corrected for voxel-level whole-brain family-wise error [FWE] based on permutation). VOI representation was additionally found in lateral prefrontal cortex (middle frontal gyrus; MFG), right superior frontal gyrus (SFG), posterior cingulate cortex (PCC), right angular gyrus, and cerebellum (Fig. 4a). Striatum and VMPFC receive dopaminergic inputs and are the two regions that are the most associated with valuation in fMRI literature. Indeed, we found that lottery’s EU was represented in striatum during the presentation of lottery (*p* < .05, corrected for whole-brain FWE; Fig. 4a), and it was overlapped with VOI cluster, suggesting the involvement of traditional valuation processing in VOI.

**Fig. 4.**
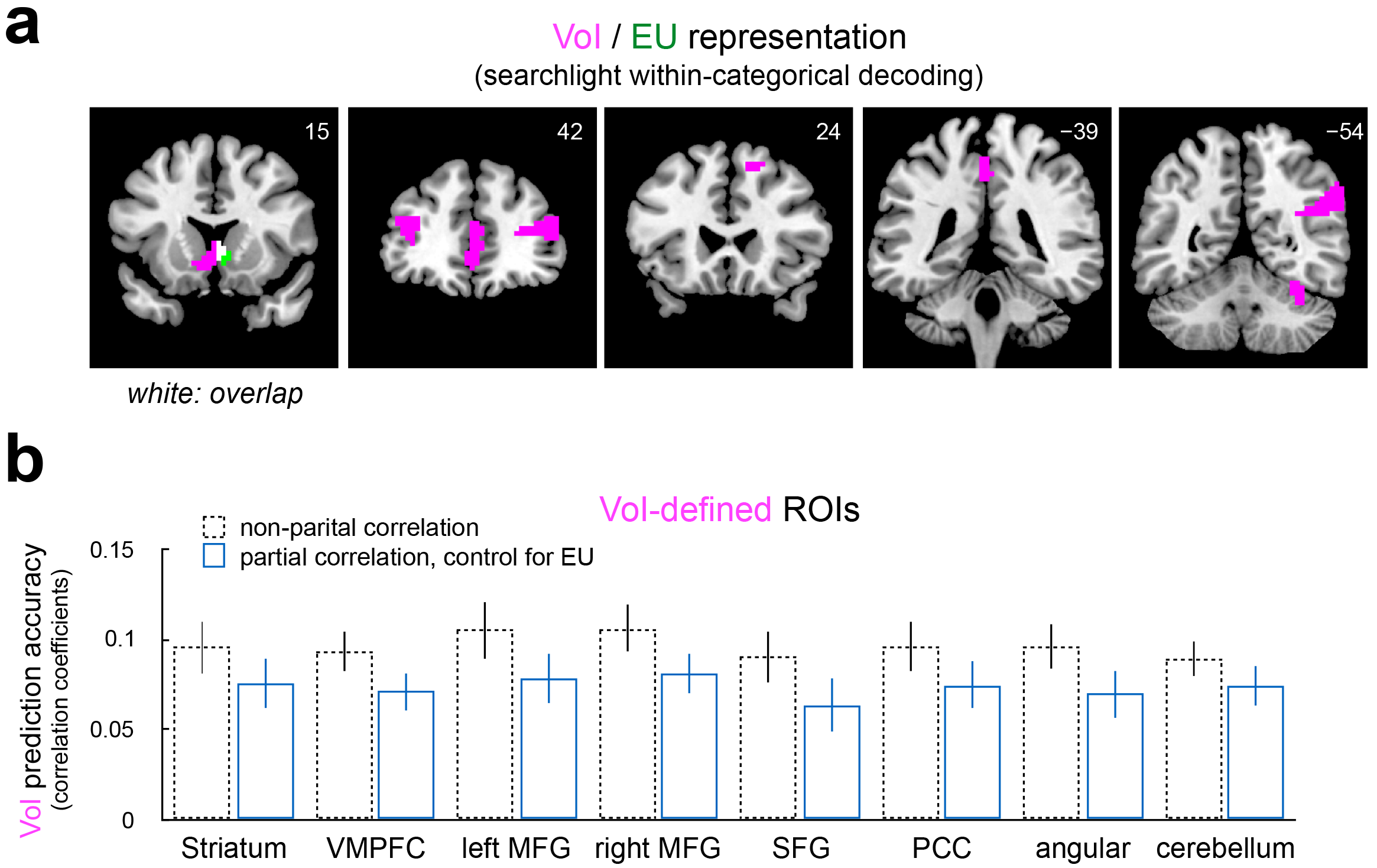
Neural representation of VOI and EU. (**a**) VOI representation was revealed in regions including striatum and VMPFC by within-categorical decoding (magenta; voxel-wise *p* < .05, whole-brain FWE corrected). EU representation was also found in striatum (green), overlapping with VOI (white). In VOI decoding, prediction accuracy was assessed while controlling for diagnosticity (see main text). (**b**) VOI representation found in (**a**) cannot be explained by re-instantiation or maintenance of EU representation. Prediction accuracy was significantly higher than zero even when EU was controlled for (blue; all *p* < .05, Bonferroni corrected). Accuracy without controlling for diagnosticity or EU was also shown for comparison (dotted). VMPFC: ventromedial prefrontal cortex, MFG: middle frontal gyrus, SFG: superior frontal gyrus, PCC: posterior cingulate cortex.

Since VOI is correlated with the lotteries’ EU (Pearson’s *r* = 0.62), some of our VOI decoding performance might have been attributable to signals related to EU rather than VOI. Indeed, EU was decodable from striatum during the lottery presentation (*p* < .05; Fig. 4a), which was overlapped with a VOI cluster. We confirmed that this was not the case; VOI decoding accuracy in all clusters was above chance when measured when EU was controlled for (*p* < .05; Fig. 4b). This supports that these regions use not only outcomes but also information diagnosticity to calculate VOI, as normatively predicted.

### Representations of VOI and EU on common currency

Having characterized representations of VOI and EU respectively, we next investigated their relationship, and in particular whether they are represented on a common neural scale. Although we observed overlap of VOI and EU clusters, this is not a strong evidence for a common scale, because these representations could be distinct at a more finegrained level. As a more direct test, we adopted cross-categorical decoding approach.

We hypothesized that, if EU and VOI are indeed represented on a common scale in striatum, SVR trained based on EU in striatum should be able to decode VOI (Fig. 3). To test if the obtained prediction accuracy was above chance while controlling for possible temporal dependency within trials, we obtained null-hypothesis distribution using permutation. In each iteration, outcomes and disgnosticity were shuffled over trials within each run such that dependency across runs was retained (e.g., all trials in which (*x*_1_, *x*_2_, π) = ($12, −$6, 2/3) were treated as if it were ($6, −$9, 1)), and trial-wise VOI labels and EU labels were given to each trial accordingly. Group-level prediction accuracy (*t*-statistics) was obtained for all permutation iterations and compared against the accuracy under the ground-truth labels.

We found that the decoder trained by EU could indeed predict VOI above the chance level (*p* < .05; Fig. 5a). This holds when information diagnosticity was controlled for in evaluating prediction accuracy, and more critically, even when EU under *s*_0_ was controlled for. This provides a clear evidence that striatum did not just maintain or reactivate EU representation; rather, it flexibly switched the content of representation within each trial from EU and VOI, presumably in preparation for the respective upcoming choices.

**Fig. 5.**
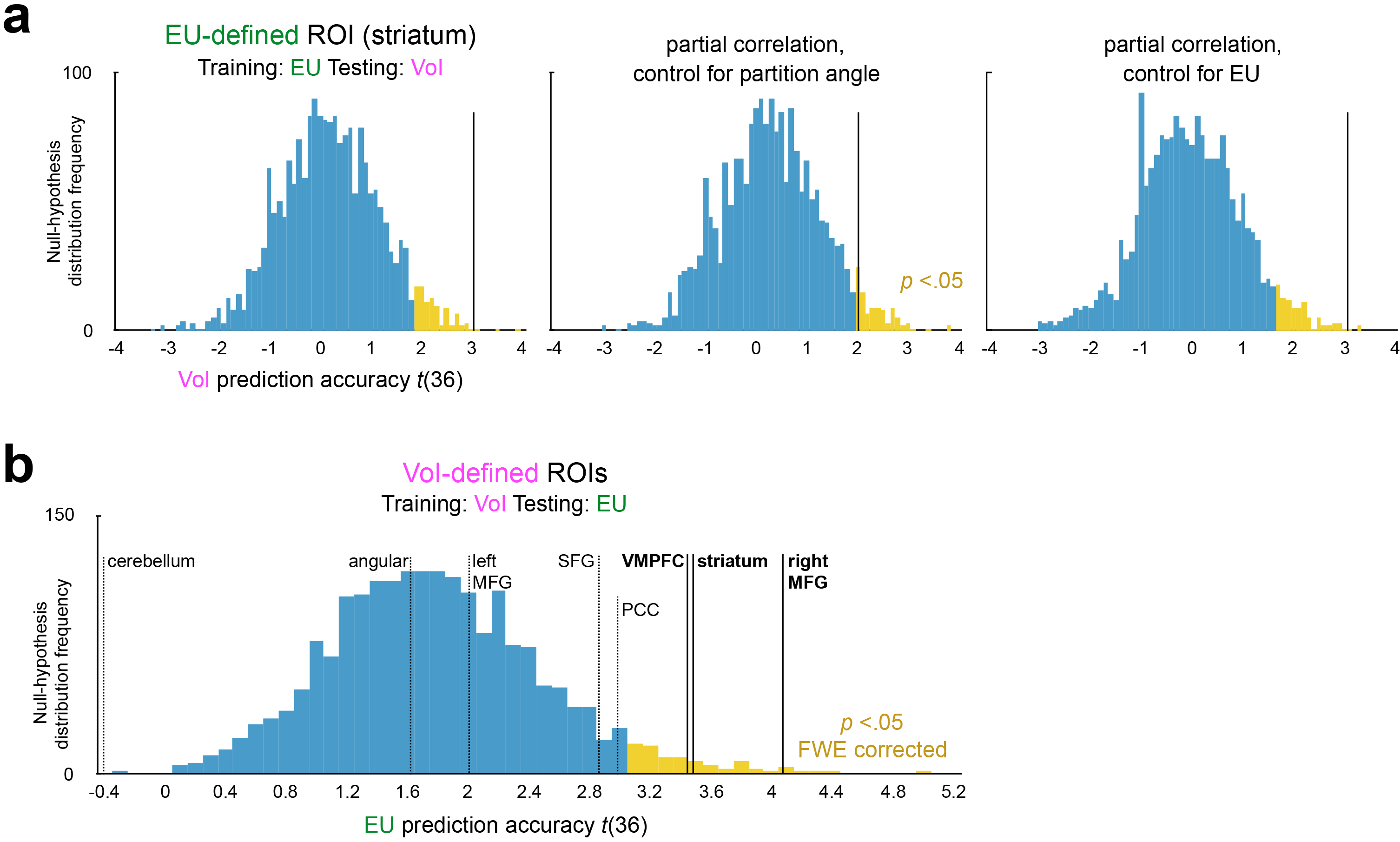
Evidence for neural common currency. (**a**) In striatum (green in Fig. 4a), decoders trained on EU predicted VOI. Shown are permutation-based null-hypothesis distribution of *t*-statistics of prediction accuracy (black vertical line: true accuracy). *Left*: no control, *middle*: controlling for diagnosticity, *right*: controlling for EU. (**b**) In striatum, VMPFC, and right MFG (magenta in Fig. 4a), decoders trained on VOI predicted EU. To control for familywise error, the null-hypothesis distribution was made of the highest accuracy among ROIs for each permutation iteration.

Lastly, to seek for further evidence for common neural code, we examined if decoders trained by VOI could be used to decode EU. To control for FWE over 8 VOI clusters reported above, we constructed null-hypothesis distribution based on the highest accuracy (*t*-statistics) over ROIs in each permutation iteration. As a result, EU prediction accuracy was deemed to be above chance in striatum, VMPFC, and right MFG (*p* < .05, Fig. 5b). Although EU was not decodable from VMPFC and right MFG in the within-categorical decoding analysis above, it may be because we had used more stringent statistical threshold. Together, these results show that human brains use a common scale to represent VOI and EU.

## Discussion

A substantial portion of our daily actions pertains to information acquisition. Particularly in a digital age where a tremendous amount of information is available at our fingertips, acquiring relevant information to an appropriate degree is as important as making use of acquired information. Going back at least to Berlyne^3^, psychologists studying functions, causes, and consequences of motivation and interests have hypothesized the relationship between exploratory and information-seeking behavior and reward system. However, only recently have researchers begun to elucidate the neural basis of adaptive information acquisition.

In their influential proposal, Kakade and Dayan^15^ hypothesized that dopamine reward system produces information seeking by multiplexing signal on extrinsic reward and some utility bonus and encouraging exploration. Consistent with this, putative non-instrumental motives, such as self-reported curiosity or stimulus novelty, correlate with BOLD signals in striatum and dopaminergic midbrain regions in humans ^9–11,32–34^. More direct evidence for dopaminergic responses to information was provided by Bromberg-Martin and Hikosaka^16^, who found that monkeys preferred advanced information on the amount of future reward, and that midbrain dopamine neurons encoded information in a highly consistent manner with reward prediction error. However, existing studies have been limited to non-instrumental information cases, and it remains unclear to what extent dopaminergic reward system is involved in adaptive, forward-looking information acquisition.

If information acquisition is indeed driven by dopamine reward system in general, we should expect two features from dopaminergic responses; first, they should be scaled according to subjective preference for information, even when it is sensitive to instrumental benefit, and second, they should be on a common scale with signals on extrinsic reward. Our results on the common valuation scale in BOLD from striatum and VMPFC are highly consistent with these predictions, because they receive massive dopaminergic projection^35^ and represent various kinds of values^36,37^, with some evidence for common currency^21,23,24,26^. In particular, our findings expand existing knowledge by showing that striatum also represents forward-looking instrumental benefits. Furthermore, our cross-categorical decoding approach provides a more direct evidence for neural common currency above and beyond regional overlaps as typically reported in brain mapping studies. Our results also predict that, when monkeys act on forward-looking instrumental benefit of information, it may also be encoded by their midbrain dopamine neurons.

We found VOI representation in other areas as well, but evidence for neural common currency was not found in most of them; cross-categorical decoding was successful in right MFG but not in the other clusters. As VOI computation requires the simulation of agents’ own choices and outcomes under possible informational states, it may need more neurocognitive recourses than other valuation, particularly working memory and planning. Relatedly, although encoding of non-instrumental information value was reported in orbitofrontal cortex (OFC) of monkeys, contrary to midbrain dopaminergic areas, it was distinct from reward encoding^38^. This suggests that, while OFC neurons may encode signals relevant to information valuation, they seem not to use a common code with other types of values^39,40^. Taken together, information valuation may be supported by widespread neurocognitive resources, and it may converge with other values for the first time in striatum and/or VMPFC. Downstream processes, such as action selection, may be largely similar whether they involve information acquisition or not.

Our study also provides insight on cognitive processes underlying information acquisition, and in particular the importance of valuation systems. We behaviorally identified at least two motives, forward-looking instrumental benefit and anticipatory utility. Other models on non-instrumental motives that are independent of reward value of outcomes, such as constant utility bonus^5^, were insufficient in explaining the observed behavior. Particularly, consistent with the notion of savoring, we found stake-dependent over-purchase of information. Our results extend the findings from the past studies on anticipatory utility, which have focused mostly on non-instrumental information and not quantitatively captured concurrent contribution of instrumental and anticipatory value for information^29,30,41^.

The possibility that anticipatory utility is an important component of motivation to acquire information opens up several important questions. One particular issue concerns the effect of dread, or utility of anticipating negative outcomes^42,43^. The effect of dread may be large enough for some people to avoid potentially negative information even when its instrumental benefit is critical, such as medical conditions^44–46^, but its relative contribution in instrumental information settings is yet to be empirically quantified. Our study could not measure its effect reliably because our subjects could reject unfavorable lotteries. Second, anticipatory utility provides a possible explanation for the phenomenon of ambiguity aversion. Intuitively, the desire for information may be causally linked to aversion to the lack thereof^13,47,48^. It may thus be not a coincidence that non-linearity of the aggregator function that determines second-order utility, a critical part of recursive utility theory, is also central to some theories on ambiguity and compound lotteries^49–51^. Our paradigm could be used to quantify anticipatory utility at the individual level and correlate with ambiguity attitude.

More generally, little is understood on how humans adopt different strategies on information acquisition under various situations, from stable to dynamic environments, and from short to long temporal horizons^1,4,22^. Although we found little support for utility bonus accounts in our experimental paradigm, it is entirely possible that they are appropriate description of exploratory behavior in more dynamic settings with longer temporal horizon^5,52,53^. Similarly, other proposed motives we did not study here, such as novelty, complexity, or surprise^1,3,8,54–57^, might be necessary or more suited to ensure adequate degree of exploration in certain circumstances, particularly outside value-based decision-making domains. While our findings on anticipatory utility cannot be explained by these concepts, it is yet to be quantified when and how much these motives contribute to behavior. This issue is interesting particularly under the light of recent progress in AI, in which agents who explore the environment based on prediction error alone can acquire knowledge to an appropriate degree and generalize it to novel situations^58^. Various potential motives have been long studied separately in respective fields, and the current study marks an important step, both theoretically and empirically, towards integrative understanding.

## Methods

### Subjects and procedure

47 healthy, naïve subjects were recruited from a subject pool at Virginia Tech. Ten subjects were removed due to excessive motion, leaving 37 subjects included in analyses (21 females, age range = 19-38, average 24.5). They were screened for standard MRI contraindications, provided informed consent, received instructions, and practiced the task prior to the scanning. They used a MR-compatible button box to interact with the task. The experiment program was on Matlab (Mathworks) and Psychtoolbox^59,60^ on a Windows PC. All protocols were approved by UC Berkeley Committee for the Protection of Human Subjects and Virginia Tech Institutional Review Board.

### Experiment task design

The task procedure is illustrated in Fig. 1a. In each trial, a lottery with two outcomes was presented as a roulette wheel, partitioned by a vertical white line at the middle (3 seconds). Ten lotteries were used: (+$12, −$9), (+$9, −$12), (+$9, −$9), (+$12, −$6), (+$6, −$12), (+$9, −$6), (+$6, −$9), (+$6, −$6), (+$12, −$3), and (+$3, −$12). After a fixation screen with variable duration (1–4 seconds), subjects chose whether to play the lottery or not (within 2 seconds). After the presentation of the same lottery (1–4 seconds), the information was presented as a magenta partition on the wheel (3 seconds). The partition was slanted by either 0° (vertical), 30°, 60°, 120°, or 150°. 0° corresponds to diagnosticity π of 1, 30° and 150° to 5/6, and 60° and 120° to 2/3, respectively. After another fixation screen with variable duration (1–4 seconds), the information cost was presented, and subjects chose whether to purchase it or not (within 2 seconds).

Scanning consisted of five runs, each comprised of 30 trials in a randomized order (150 trials in total). Each combination of lotteries and diagnosticity was shown once in each scanning run (10 lotteries × 3 levels of diagnosticity). The information cost was either ¢5 (3 trials per run), $1 (8 trials), $2 (8 trials), $3 (8 trials), or $9 (3 trials), and varied independently from lotteries and diagnosticity. Trials in which subjects did not respond within 2 seconds were discarded from the behavioral and fMRI analysis.

After the scanning, five trials were randomly selected and implemented into the actual monetary payment. If they had accepted a selected lottery, a green dot appeared on the wheel’s perimeter, which determined the outcome (gain or loss); if they had rejected it, subjects did not receive gain or loss. If subjects had not purchased the information, the green dot’s location followed uniform distribution across the perimeter (i.e., the two possible outcomes were equally likely). If subjects had purchased the information, one side of the magenta partition was brightened, indicating that the dot’s location followed uniform distribution within the brightened side. They could change their original choice on the lottery (accept or reject) accordingly. The brighter side was chosen randomly. The outcome from the five selected lotteries were the averaged and added to the baseline payment for completion.

### Behavioral modeling

Unless noted otherwise, free parameters were estimated using the standard maximum likelihood estimation procedure (mle function on Matlab).

First, subjects’ group-level utility function was estimated from their choices on lotteries during scanning. It involved four free parameters: power on the positive and negative domains, a multiplicative term on the negative domain, and the temperature parameter of soft-max binary choice function. The estimated utility function was *u*(*x*) = *x*^0.68^ (*x* > 0), −1.74 · (−x)^0.49^(*x* < 0), indicating strong risk and loss aversion.

Second, trial-wise prediction of standard VOI was derived as a function of lottery outcomes (*x*_1_ and *x*_2_), information diagnosticity (π), and the estimated utility function *u*(.). A fully deterministic, EU-maximizing (i.e., greedy) policy was assumed. VOI was obtained as the sunk cost at which greedy agents are indifferent to information purchase. Therefore, VOI is obtained as the cost *c* that satisfies

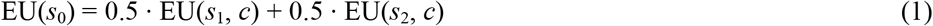

where

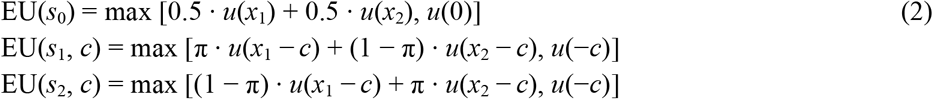

VOI was numerically obtained by minimizing the absolute difference between LHS and RHS in Equation 1 (fminsearch function on Matlab).

Third, subjects’ trial-wise information purchase choices were modeled by standard VOI (Fig. 2b). This involved only one free parameter: the temperature of the soft-max choice function that maps VOI minus the cost onto binary choices. The goodness-of-fit of this model was evaluated as log likelihood of all choices (summed over subjects) and statistically tested by permutation, in which null-hypothesis distribution was obtained by shuffling the labels of the lottery outcomes (*x*_1_ and *x*_2_) and the diagnosticity (π) for 2,000 times and compared to the ground-truth model fit.

Fourth, standard VOI modeling was compared against the constant bonus account and the entropy bonus account. This involved two parameters: the magnitude of bonus (constant term or reduction in entropy, log(0.5) − πlog(π) − (1 − π)log(1 − π), respectively) and the temperature parameter of soft-max function. To statistically compare their log likelihoods to standard VOI model (Fig. 2c), 10-run, 10-fold cross validation across subjects was conducted. The whole data was randomly split into 10 datasets (3 or 4 subjects each), and bonus parameters were estimated from 9 datasets and evaluated on the hold-out dataset (soft max temperature was estimated anew in the hold-out dataset). This procedure was repeated for each hold-out dataset, and then for 10 dataset splits. This yielded 100 log-likelihood values. They were compared against log likelihood of standard VOI model on the same hold-out dataset using *t*-test. Degree-of-freedom of *t*-test was set to 10 instead of 99 to correct for dependency among iterations^61^.

Fifth, the composite models of standard VOI and utility bonus accounts (constant bonus or entropy reduction bonus) were tested (Fig. 2f). Two free parameters were estimated: the magnitude of bonus and temperature of soft-max function that mapped [VOI plus bonus minus cost] onto binary choices. Their goodness-of-fit was compared against standard VOI model using 10-run, 10-fold cross validation, as described above.

Sixth, and lastly, the generalized VOI under recursive utility was tested (Fig. 2e). In this model, equations (1) and (2) are modified as

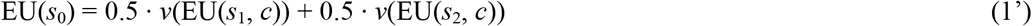

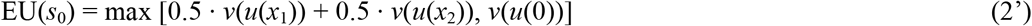

where *v*(.) is the aggregator function that returns second-order utility. *v*(.) included three free parameters: power on the positive domain, power on the negative domain, and a multiplicative term on the negative domain. *v*(.) and the temperature of the soft-max choice function. Here, because the solution of (1’) has non-linear dependence on *v*(.), this is a two-level estimation problem: we need to find *v*(.), under which estimated VOIs achieve the lowest log likelihood of binary choices. Thus, instead of mle function, we estimated *v*(.) using Nelder-Mead simplex algorithm^62^ using a custom-made Matlab script. Comparison to alternative models (Fig. 2f) were carried out using 10-run, 10-fold cross validation, as described above.

We also evaluated stake-dependent deviations from the instrumental-only and generalized VOI predictions (Fig. 2b, e). For each trial in each subject, deterministic trial-wise prediction (purchase or forgo) was obtained using best-fit parameters. Next, trials were separated into two categories according to lotteries’ EUs under *s*_0_ (median split of 10 lotteries). Difference in the predicted and actual purchase probability in each category was then averaged across subjects.

### MRI acquisition

MR images were acquired by a 3T Siemens Trio scanner and a 12-channel head coil. Prior to the task, T1-weighted structural images (1mm × 1mm × 1mm) were obtained using magnetization-prepared rapid-acquisition gradient-echo (MPRAGE) pulse sequence. During the task, functional images were obtained using T2*-weighted gradient-echo echo-planar imaging (EPI) pulse sequence (TR = 2000ms, TE = 30ms, voxel size = 3mm × 3mm × 3mm, inter-slice gap = 0.3mm, in-plane resolution = 64 × 64, 32 oblique axial slices). Slices were tilted by approximately 30 degrees from AC-PC line to reduce signal dropout from orbitofrontal cortex^63^.

### MRI preprocessing

Motion correction and slice-time correction were applied to EPI images on SPM12 (http://www.fil.ion.ucl.ac.uk/spm/). Neither smoothing or normalization was applied as a part of preprocessing, but after decoding analysis (see below).

#### MRI analysis

##### General linear modeling (GLM)

We used voxel-wise, trial-wise activation estimates as features in support vector regression (SVR). They were obtained by GLM on SPM12. Each of the five runs for each subject was modeled separately. Each GLM included one regressor modeling lottery presentation (3-second boxcar) and one regressor modeling information presentation (3-second boxcar) for each trial with responses. Additional regressors modeled these epochs in all trials without responses (excluded from the following SVR). All button presses were modeled by two additional regressors, one for right-hand responses and one for left-hand responses. All of these event-related regressors were convolved with the SPM’s double-gamma canonical hemodynamic response function. In addition, GLMs included movement parameters estimated in the motion-correction procedure, 128-sec high-pass filtering, and AR(1) model of serial autocorrelation.

##### Within-categorical decoding

Our approach is schematically illustrated in Fig. 3. The decoding analysis used custom-made Matlab scripts and adopted linear kernel epsilon-SVR in LIBSVM package^64^. Cost parameter and epsilon parameter were pre-set at 1 and $0.1, respectively.

Within-categorical decoding with searchlight was conducted to look for neural representation of VOI and EU (Fig. 4a). It consisted of two steps, individual-level and group-level. In individual-level analysis, we evaluated SVR’s decoding accuracy of VOI during information presentation and that of EU during lottery presentation. Five-fold leave-one-run-out cross validation was conducted for each spherical searchlight (radius: 10 mm). In each fold, a decoder was trained on features (trial-wise activation pattern estimated in GLM above) from four runs (up to 120 trials) so that it finds a linear model that predicts value labels. The trained decoder was then used to predict value labels in the hold-out run. Regarding value labels, we used trial-wise VOI estimated under the recursive utility model with best-fit parameters and trial-wise EU obtained based on estimated group-level utility function (see above). Due to the design of the task, trial-wise VOI was highly correlated with information diagnosticity π. In order to find clusters which activation was associated with VOI above and beyond π, we measured VOI decoder’s prediction accuracy as Pearson partial correlation between predicted and actual VOIs, controlling for π. EU decoder’s prediction accuracy was measured as (non-partial) correlation between predicted and actual EUs. Measured accuracy was then z-transformed, averaged over the five cross-validation folds, and assigned to the central voxel of the searchlight. This created subject-wise prediction accuracy maps, one for VOI and one for EU.

For group-level analysis, subject-wise accuracy maps were normalized to MNI template using SPM12’s DARTEL procedure. DARTEL consists of two steps: non-linear transformation to the average brain among subjects and affine transformation to the template. Normalized maps in MNI space were then smoothed with a Gaussian kernel (8-mm FWHM). Group-level statistical significance was evaluated using random-effect model as conventional SPM analyses. We used voxel-level threshold *p* < .05, corrected for whole-brain family-wise error (FWE). FWE correction was conducted based on non-parametric permutation using SnPM13 package^65^.

In an additional ROI analysis (Fig. 4b), we examined if decoding accuracy could be explained by EU representation instead of VOI. ROIs were defined at *p* < .05 (voxel-level, FWE corrected) and transferred to subject-wise local spaces using the reverse transformation of DARTEL. Next, prediction accuracy of VOI decoder was tested in searchlights centered on every voxel within each ROI, while controlling for EU (instead of π). As a reference, prediction accuracy without any control (non-partial Pearson correlation) was also measured. Accuracy measures were then averaged within each ROI, and then across subjects.

##### Cross-categorical decoding

Within-categorical decoding analysis revealed a cluster in striatum from which EU was decodable, and eight clusters from which VOI was decodable. We next conducted cross-categorical decoding to ask if they were represented on a neural common currency. As above, ROIs were defined at *p* < .05 (voxel-level, FWE corrected) and transferred back to local spaces. Prediction accuracy was averaged across searchlights centered on voxels within each ROI.

We first examined if decoders trained on EU could predict VOI (Fig. 5a). Decoders trained in within-categorical decoding were used, but they were applied to activity pattern in information presentation epoch to predict VOI (Fig. 4). As before, we evaluated VOI prediction accuracy in three ways: non-partial correlation, partial correlation controlling for diagnosticity, and partial correlation controlling for EU. The last control is particularly important as it addresses the concern that striatum might re-instantiate (or maintain) EU representation over time. Group-level *t*-statistics was then obtained over these measures averaged within the ROI.

To test if the obtained prediction accuracy was higher than chance, it was important to control for possible signal dependency within trials (e.g., signals in information presentation epoch could be dependent on signals in lottery presentation epoch due to autocorrelation unexplained in GLM). To do so, we conducted permutation with 1,000 iterations. In each iteration, we shuffled value labels over trials within each run, but retained the dependency between VOI labels and EU labels, as well as dependency across runs. Prediction accuracy measures were then obtained for each iteration exactly as the original analysis (fivefold, leave-one-run-out cross validation). Fig. 5a shows the histogram of *t*-statistics under permutated labels (i.e., null-hypothesis distribution).

Next, we examined if decoders trained on VOI could predict EU (Fig. 5b). The procedure of this analysis (VOI to EU) was overall the same as above (EU to VOI). In order to control for family-wise error among 8 VOI ROIs, null-hypothesis distribution of *t*-statistics was formed by the highest *t*-statistics among 8 ROIs in each permutation iteration (1,000 iterations in total). As a result, we obtained the null-hypothesis distribution which center was shifted positively from zero.

